# Summation of contrast across the visual field: a common “fourth root” rule holds from the fovea to the periphery

**DOI:** 10.1101/2025.05.06.649158

**Authors:** Alex S Baldwin, Tim S Meese

## Abstract

Increasing the area of grating-like stimuli reduces their contrast detection thresholds. Characterising the visual system’s summation rule this way provides insights into early visual architecture. Previous work in the fovea has found linear summation over short distances, consistent with integration within the receptive fields of early cortical neurons. Beyond this range, the benefit of stimulus area is reduced. Here, we investigated whether the same integration rule holds for stimulus elongations centred at different positions across the visual field. We did this for “tiger tail” strips of grating (growing orthogonally to the major axis of the early receptive fields) in the fovea, parafovea (3 deg), and periphery (10.5 deg). The interpretation of results from previous studies has been complicated by variation in local contrast sensitivity across the visual field. We addressed this here by using detailed maps of the inhomogeneity for each participant (their “witch hat”) to generate “compensated” stimuli where the local stimulus contrast was amplified by the reciprocal of their local sensitivity. Our results followed a common fourth-root summation rule for tiger-tails in the fovea, parafovea, and periphery. We explained this by a “noisy energy” model that combined: i) a “witch hat” sensitivity surface, ii) linear filtering by receptive fields, iii) square-law contrast transduction, and iv) an internal template to direct the observer’s attention to the spatial extent of the stimulus. Fitting this model with a single global sensitivity parameter accounted for foveal and parafoveal results (56 thresholds), with one further parameter needed to model the periphery (84 thresholds).

## 1 Introduction

### 1.1 Summation of contrast to threshold for stimuli of different sizes

Area summation studies investigate how the visual system combines signals over space. For example, one can measure the relationship between the area of a sinusoidal grating and its contrast detection threshold (the lowest contrast at which a stimulus is detected with criterion probability). The Michelson contrast of a stimulus is defined as

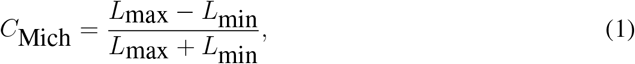

and is proportional to the square root of the power of the sinusoid that renders the grating stimulus.

Under Signal Detection Theory (SDT; Green and Swets, 1966), the ability to detect low-contrast targets is limited by intrinsic noise in the visual system (Gregory and Cane, 1955). In general, larger gratings can be detected at lower Michelson contrast. This produces a negative slope in a log-log plot of contrast threshold against stimulus area, where the steepness of the slope provides information about the summation process. In this article, we plot contrast thresholds expressed in logarithmic dB units on the y-axis

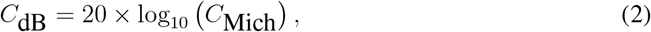

and, for ease of comparison, we apply the same transformation to the stimulus area for the x-axis.

When plotting threshold against area, a slope of −1 is the signature of a linear combination of contrast across the visual field (provided the detection-limiting noise remains constant as the stimulus size increases). For linear summation, performance for detecting a small grating with a Michelson contrast of 2% would be the same as that for detecting a grating with twice the area at a contrast of 1%. Summation slopes that are shallower than this indicate nonlinear characteristics of the detection process. For example, if the threshold were determined by the contrast energy of the stimulus (which is proportional to the contrast power integrated over space) then the summation slope (on double-log axes) would be −½ (Manahilov and Simpson, 1999).

### 1.2 The probability summation account of area summation to threshold

Empirical results from previous psychophysical studies have found summation slopes of approximately −¼ (“fourth root” summation; Bonneh and Sagi, 1999). These have often been interpreted as probability summation (Robson and Graham, 1981) where each part of a strip of grating is detected by independent local mechanisms. Because the number of viable detecting mechanisms increases with the area of the grating, there is a greater probability that at least one mechanism will detect the target (Sachs et al., 1971). Models of the benefit to sensitivity from probability summation were originally formulated under High Threshold Theory (Green and Swets, 1966), where local mechanisms have a binary response (“detected” or not “detected”). In this case, a summation slope prediction (−1/*β*) can be derived from the slope (*β*) parameter of a Weibull psychometric function (Quick, 1974; Robson and Graham, 1981). The psychometric slope characterises how the probability for detection changes with signal strength. Under High Threshold Theory, when *β* = 4 the probability summation prediction corresponds with fourth root summation.

However, it has been demonstrated that psychophysical behaviour is inconsistent with the binary High Threshold Theory framework (Swets, 1961a,b; Corso, 1963; Nachmias, 1981). Instead, results suggest that observers have access to a continuous response from each local mechanism, subject to additive noise. Consider an *M*-interval forced-choice task, where the observer must indicate in which of *M* temporal intervals a target was presented. Under SDT, if the target stimulates a single visual mechanism, such that the task is a decision over *M* responses, the decision can be made through a max operation where the single greatest response indicates the presence of the target (Tanner and Swets, 1954). This can be extended to develop a model of spatial summation. When multiple mechanisms are available to detect the target, a maximum response can also be taken over space in each interval. This means the decision is based on the single greatest activation that occurs in each trial. Under plausible assumptions, this model also predicts fourth root summation with stimulus area (Tyler and Chen, 2000; Meese and Summers, 2012; Kingdom et al., 2015). For convenience, we will continue to refer to this idea as “probability summation”, though the term is a misnomer under SDT.

### 1.3 Evidence for a noisy energy model

Our recent work on area summation has favoured a “noisy energy” model over probability summation (Meese and Summers, 2007; Meese, 2010; Meese and Summers, 2012; Baldwin and Meese, 2015). In this model, the responses of local contrast-driven mechanisms are squared and summed within a template that roughly corresponds with the size and location of the stimulus. The size of the template plays a double role in influencing thresholds; not only does it set the integration range for the signal, but the response at each location is also perturbed by independent local noise. The variance of the noise that is integrated (to form the denominator in the signal-to-noise ratio) therefore increases in proportion to the area of the template.

In cases where the template in the noisy energy model exactly matches the stimulus area, summation obeys a fourth root rule (Meese and Summers, 2012). This derives from a combination of: i) the square-law transduction of contrast into local mechanism responses, and ii) the quadratic effect of the integration of area-dependent noise and signal. For more complex stimuli the template-matching process breaks down (Meese and Summers, 2007; Baker and Meese, 2011; Meese and Summers, 2012; Baldwin and Meese, 2015). For example, an experiment using a “Battenberg” checkerboard pattern of signal regions (containing luminance contrast) and non-signal regions (which were blank) revealed a potent square root (quadratic) summation process (Meese, 2010). This was explained by proposing a limitation in the template profile such that it matched the overall spatial extent of the signal, but not the spatial modulations of local contrast (i.e. the template did not accommodate the non-signal “gaps” in the Battenberg stimulus). With this arrangement, the spatial integration of additive internal noise does not change when the Battenberg “gaps” are filled with signal, leaving only the quadratic effect of square-law signal transduction. This quadratic result (Meese, 2010) was critical in deciding between the noisy energy model and probability summation models which typically produce fourth root summation (or thereabouts), as described above (see also Meese and Summers, 2012, for further evidence from exploiting model constraints).

### 1.4 Effects of the visual field inhomogeneity in local contrast sensitivity

Most area summation experiments for periodic spatially band-pass stimuli^2^ fall into one of two categories. In the first, contrast detection thresholds are measured for centrally-presented gratings for a wide range of grating areas (e.g. Howell and Hess, 1978; Rovamo et al., 1994; Meese and Summers, 2012). However, the summation slopes in these experiments are confounded by the inhomogeneity in contrast sensitivity across the visual field (Hilz and Cavonius, 1974). As the size of the grating increases, it extends into more eccentric regions of the visual field where contrast sensitivity is lower. The decline in log-sensitivity with eccentricity was classically understood to be linear, with each unit of translation toward the periphery reducing sensitivity by a constant factor (Robson and Graham, 1981). The decline can be modelled as scale-invariant (having a common slope when eccentricity is given in periods of the grating sinusoid, or “carrier cycles”) for a broad range of spatial frequencies (e.g. 1.6 to 12.8 c/deg in Pointer and Hess, 1989). The scale-dependent decline in sensitivity (in absolute units of eccentricity such as degrees of visual angle) determines the spatial resolution at each eccentricity (Watson, 2018).

In Baldwin et al. (2012), we mapped the variation in sensitivity across the central visual field with finer sampling than that used in previous studies. We found the decline in log-sensitivity to be *bi*-linear rather than linear (on a plot of log-sensitivity vs linear eccentricity). An initial steep decline was followed by a shallower slope (having from half to a quarter of the initial gradient) with the inflection point at a fixed number of grating periods (and therefore scale-invariant). The gradients of the two slopes (steep and shallow) varied with the polar angle of the spatial trajectory along which testing was performed, consistent with the Horizontal-Vertical Anisotropy and Vertical Meridian Asymmetry reported by Abrams et al. (2012).

The measurements made in Baldwin et al. (2012) provided the basis for personalised “Witch Hat attenuation surfaces” of contrast sensitivity, and were used in spatial summation experiments by Baldwin and Meese (2015) to counteract spatial inhomogeneity by multiplying target gratings of various sizes by the inverse of the surface. This allowed a cleaner measure of area summation by factoring out the confounding influence of the inhomogeneity in local sensitivity.

In the second of the two types of area summation study, smaller strips of grating are used to investigate local summation behaviour and the effects of stimulus aspect ratio (Robson and Graham, 1981; Mayer and Tyler, 1986; Polat and Norcia, 1998; Manahilov et al., 2001; Foley et al., 2007; Meese and Hess, 2007). These strips of grating are often presented to regions of the visual field outside the fovea where contrast sensitivity is more uniform. These strips can “grow” in size in a direction either parallel (“skunk tails” in Meese and Hess, 2007) or perpendicular (“tiger tails”) to the bars of the grating. There is evidence for summation along skunk tail stimuli involving additional mechanisms (Chen and Tyler, 1999; Chen et al., 2019, 2023), perhaps related to the integration of contours (Field et al., 1993). In the study here, we are primarily concerned with summation along the width of our tiger tail stimuli (orthogonal to the grating stripes), though our use of stimuli with different heights does allow some examination of summation along the grating stripes.

For very small stimuli with an area of less than one square cycle (Foley et al., 2007) the summation slopes are steep. This is thought to reflect linear summation within the footprint of early receptive fields (Chen and Tyler, 1999; Meese, 2010). Slightly larger stimuli presented to the periphery, with an area of less than 32 square cycles, have log-log summation slopes around −½ (Manahilov et al., 2001; Meese and Hess, 2007). For even larger stimuli, the slopes are around −¼ (Robson and Graham, 1981). Summation slopes for foveal stimuli are typically flatter due to the inhomogeneity in contrast sensitivity across the visual field. If this is not compensated, then as the stimulus “grows” in size, it encroaches progressively less sensitive regions of the visual field, and the benefit of signal area diminishes (Baldwin et al., 2012).

### 1.5 Does area summation obey a single rule across the visual field?

Our goal here was to further our understanding of early spatial vision by investigating whether a single rule could explain spatial summation for stimuli centred at various locations across the visual field (i.e. in the fovea, the parafovea, and the periphery). In the studies discussed above, summation was typically investigated over a relatively narrow range of stimulus sizes and, in many cases, at just a single location in the visual field. One approach to compare performance at different locations in the visual field has been to scale stimuli according to a “Cortical Magnification Factor” (Cowey and Rolls, 1974; Strasburger et al., 2011). For example, scaling the spatial frequency and area of peripheral grating stimuli by the inverse of the magnification factor (normalising contrast sensitivity within *±* a factor of two in Rovamo and Virsu, 1979; Jigo et al., 2023). In this study however, we are interested in comparing the detection of stimuli with the same spatial scale at different locations in the visual field.

For effective investigation of spatial summation, each part of the stimulus should be equally detectable. Previous work achieved this by testing across regions of peripheral vision where contrast sensitivity was shown to vary only slightly (Robson and Graham, 1981). Our “Witch Hat” compensation for the inhomogeneity in contrast sensitivity (Baldwin and Meese, 2015) improved on this by “flattening” the effective contrast of stimuli across the foveal and parafoveal regions. We then measured thresholds for various stimulus sizes centred on the fovea, where the potential benefit of spatial integration was free of interference from inhomogenous contrast sensitivity.

In the study here, we measured area summation slopes for stimuli having a single spatial frequency (4 c/deg) and a wide range of stimulus sizes, spanning the interesting transition between steep and shallow slopes that appear to divide the conclusions across previous studies. Stimuli were presented at *three* visual field locations, with measurements made both with and without compensation for visual field inhomogeneity. Motivated by our previous work (Meese and Summers, 2012; Baldwin and Meese, 2015), we then investigated whether summation across the visual field could be accounted for by a common “fourth root” rule (consistent with our “noisy energy” model).

## 2 Methods

### 2.1 Participants

All participants were volunteers who gave their informed consent before participating in the study, which was conducted in accordance with the Declaration of Helsinki. Procedures were approved by the Ethics Committee of the School of Life and Health Sciences at Aston University (approval #856). The three participants (ASB, DHB and SAW) were 22, 28 and 46 years old respectively. Participants wore optical correction appropriate for the viewing distances tested when required. All experiments were performed binocularly with natural pupils.

### 2.2 Equipment

Stimulus presentation was performed using a CRS ViSaGe (Cambridge Research Systems, Rochester, UK), which was used with a gamma-corrected CRT monitor (Eizo Flexscan T68) to achieve 14-bit grayscale precision. The monitor had a refresh rate of 120 Hz, and a mean

### 2.3 Stimuli

Our “tiger-tail” stimuli were formed from 4 c/deg vertical “Battenberg” signal elements (Meese, 2010) arranged into rectangular strips (Figure 1). Each element was a single cycle of sinusoidal grating multiplied by an orthogonal cosine-phase half-cycle at half the target spatial frequency. An alternative stimulus design would have been to define a family of rectangular gratings of various sizes. It is generally preferable to soften the edges of grating stimuli by modulating the contrast with an envelope such as a Gaussian (to make a Gabor patch) or a raised-cosine (as in Baldwin and Meese, 2015). This complicates the definition of the stimulus extent, and consequently the calculation of the effective stimulus area. For the Battenberg stimuli used here, each micropattern element contains its own envelope. This means that the area of a 1 *×* 4 rectangular stimulus is simply four times the area of a 1 *×* 1 stimulus. Our choice of micropattern stimuli over simple gratings made no difference to the sinusoidal horizontal cross-section taken through the centre of a “row” of elements: The abutting single cycles of sinusoidal grating combined into a single continuous sinusoid. In their vertical cross-section however, the envelopes of our stimuli were rectified cosines at half the spatial frequency of the stimulus. This is accommodated in our modelling.

**Fig 1:**
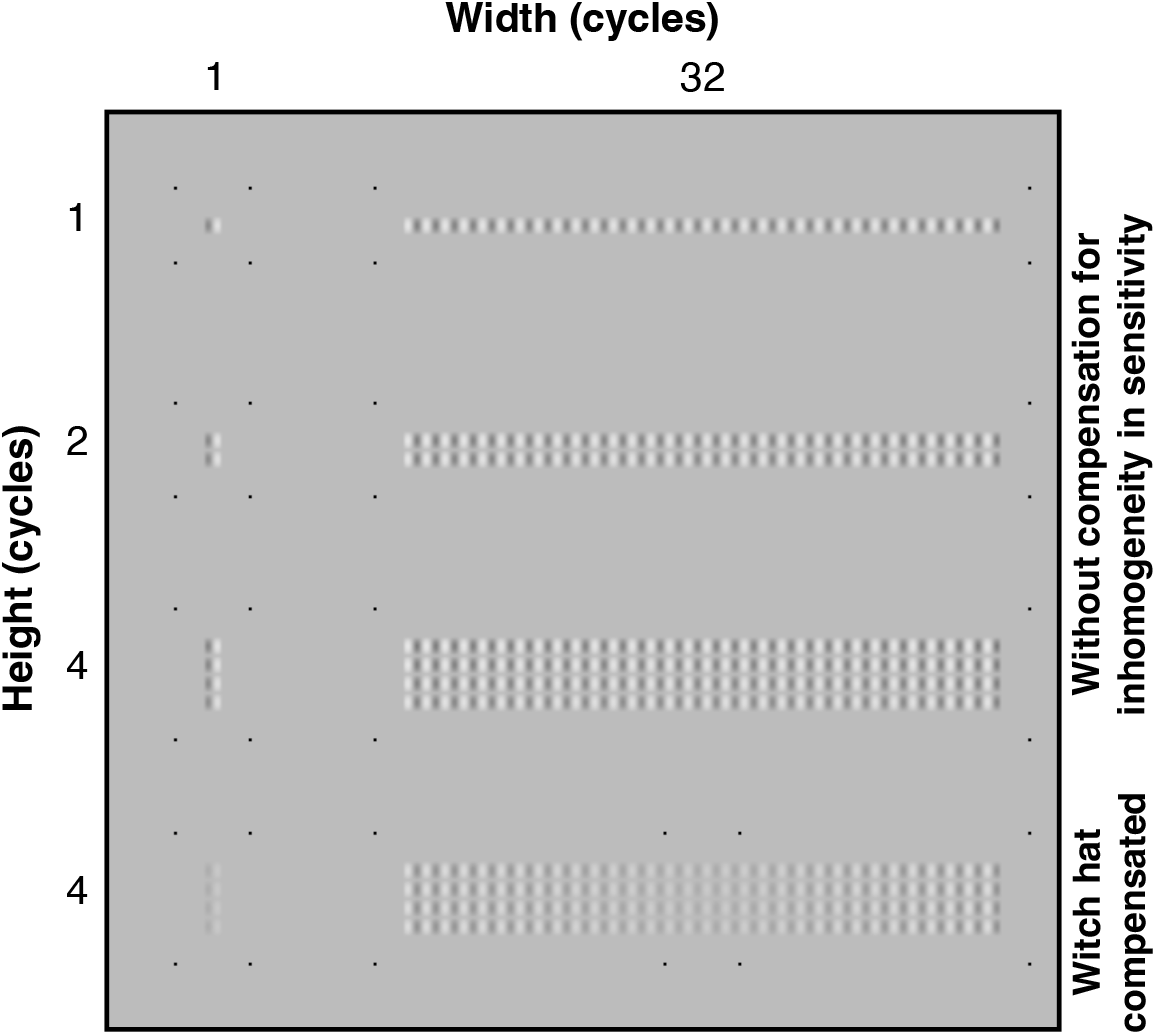
The first three rows show the narrowest (1 cycle, left) and widest (32 cycles, right) stimuli for all three heights (1, 2 and 4 cycles). These are the non-compensated stimuli, with each micro-pattern element rendered at the same contrast. The last row shows the effect of the Witch Hat compensation. Also shown are the quads of dots that served to indicate the stimulus extent to the participant. luminance of 75 cd/m^2^. The monitor was viewed from a distance of 1.2 metres. At this distance, there were 48 pixels per degree of visual angle (deg), giving the 4 c/deg stimuli used in this study 12 pixels per cycle.

Stimuli were presented with six different widths (1, 2, 4, 8, 16 and 32 cycles) and three different heights (equivalent to 1, 2 and 4 cycles). The 1 and 4 cycle high stimuli were used in all six stimulus width conditions, whereas the 2 cycle high stimuli were used only with the narrowest (1 cycle) and widest (32 cycles) stimulus widths. The nominal areas of the stimuli were calculated as the product of the width and height. These sizes were chosen to cover most of the range tested in previous studies.

Our stimulus selection included a wider range of widths than heights, focusing our investigation on “tiger tail” summation along the axis perpendicular to the stripes of the carrier grating. We supposed this would tap a simpler summation process than that for increasing height, where stimuli are grown *along* the axis of their stripes (“skunk tails”). With potential “skunk tail” effects established for each particular height (1, 2, or 4 cycles) for our narrow stimuli (1 cycle wide), orthogonal “tiger tail” summation could then be investigated by growing the stimulus width (at that height). However, including stimuli of multiple heights and widths also allowed us to compare sensitivity to stimuli with the same area but different combinations of widths and height (e.g. an area of four square cycles achieved with stimuli that are either one cycle in height and four cycles in width, or vice-versa).

Two types of stimuli were generated: those with flat contrast profiles, and those with Witch Hat compensation. The compensated stimuli were produced by multiplying the original stimuli by the inverse of the Witch Hat attenuation surface measured for each participant in Baldwin et al. (2012). This gave them an effectively flat internal signal profile after the attenuating effects of visual field inhomogeneity (Baldwin and Meese, 2015). The nominal contrast of the compensated stimuli was their contrast before the compensation was applied. Consequently, the expected threshold for these stimuli no longer depended on their locations in the visual field.

### 2.4 Procedures

The experiment was controlled using an in-house software suite (“Liberator”) written in Delphi (Borland Software Corporation, California) at Aston University. Three eccentricity conditions were tested. We chose to make our measurements along the upper vertical meridian, consistent with Robson and Graham (1981). Stimuli were presented centred either at central fixation or at two eccentric locations (both 12 and 42 stimulus carrier cycles superior to the fixation point, equivalent to 3 and 10.5 degrees of visual angle), as shown in Figure 2. We refer to these three field conditions as foveal (taken as the area within 2 deg of fixation), parafoveal (2 to 10 deg), and peripheral (beyond 10 deg), respectively. Because the stimuli extended over space, there was overlap in the eccentricities tested across the three field conditions. For the 32-cycle wide stimulus centred at fixation, the edges were at 4 deg. For parafoveal and peripheral stimuli the edges of the widest stimuli reached 5 deg and 11.2 deg respectively. Participants ASB and DHB were tested at all three eccentricities, whereas SAW was tested only for the foveal and parafoveal conditions.

**Fig 2:**
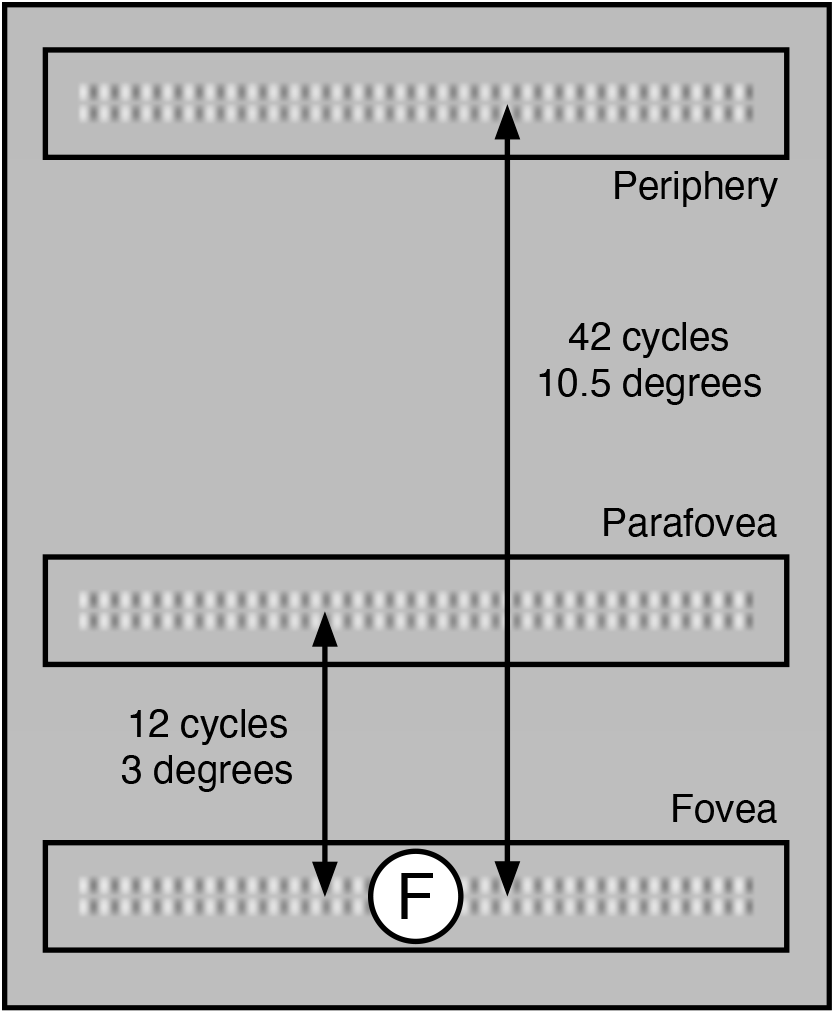
Stimulus locations used in this study (not to scale) for example stimuli two elements tall and thirty-two elements wide. The “F” icon marks the fixation location, but neither this nor a central fixation point was present in the study. Instead, the point of fixation was inferred from a quad of points around the stimulus in the foveal condition or marked directly by a 2 pixel square black dot in the parafoveal condition, or by a red LED in the peripheral condition.

The spatial extent of each stimulus was indicated by a quad of black dots surrounding its corners. For the foveal and parafoveal conditions, these dots were 2 x 2 pixel squares. In the peripheral condition, their size was increased to 4 x 4 pixels to ensure visibility. The method for marking the intended point of fixation depended on the field condition. For stimuli presented in the fovea, this was an additional quad of dots around that location (after Summers and Meese, 2009). In the parafoveal condition, we used a single 2-pixel square dot. In the peripheral condition, a dim red LED was used for fixation. This was positioned below the monitor, coplanar with the display screen, such that the distance between the LED and the centre of the stimulus was 10.5 degrees of visual angle.

Stimuli were blocked by size and location. The non-compensated and Witch Hat compensated conditions were interleaved to encourage participants to adopt the same attention strategy across the two conditions. This was motivated by our expectation that the wider non-compensated stimuli would be most visible at their centre. If this condition was tested in a separate block from the compensated stimuli (which should be equally visible across their entire span), the participant may have restricted their attentional window (Tyler and Chen, 2000) to the most salient region. The interleaving was designed to counter this potential confound. Thresholds were measured using a two-interval forced-choice three-down, one-up staircase procedure. Participants were given audible feedback as to whether they correctly chose the target interval. Each condition was repeated four times by each participant, testing the blocks in a randomised order. Contrast detection thresholds for each repetition were calculated by fitting a Weibull function to the data using Palamedes (Prins and Kingdom, 2018). Mean thresholds and standard errors (in dB) were calculated across repetitions.

## 3 Results

We first consider the results for different stimulus sizes and eccentricities before showing how the foveal and parafoveal data (56 thresholds per participant) can be modelled with a single fitted parameter per participant. Extending that to include the peripheral data (bringing us to 84 thresholds, and taking us beyond the region where we measured the Witch Hat attenuation surfaces used in our stimuli and modelling) requires one further fitted parameter per participant.

### 3.1 Width summation without compensation for visual field inhomogeneity

Contrast detection thresholds for stimuli of various widths presented to the fovea, parafovea, and periphery are shown in Figure 3, with subplots showing results from the three participants. Each data series shows a reduction in threshold as stimulus width increased but as height remained constant at either 1 or 4 cycles (see legend). More generally, thresholds decreased with increasing stimulus area across conditions. For stimuli presented to the fovea (filled symbols), summation was initially at its steepest, and then flattened out for larger stimulus sizes. We attribute this to the visual field inhomogeneity for contrast sensitivity, consistent with previous foveal results for non-compensated stimuli (e.g. Robson and Graham, 1981; Polat and Tyler, 1999).

**Fig 3:**
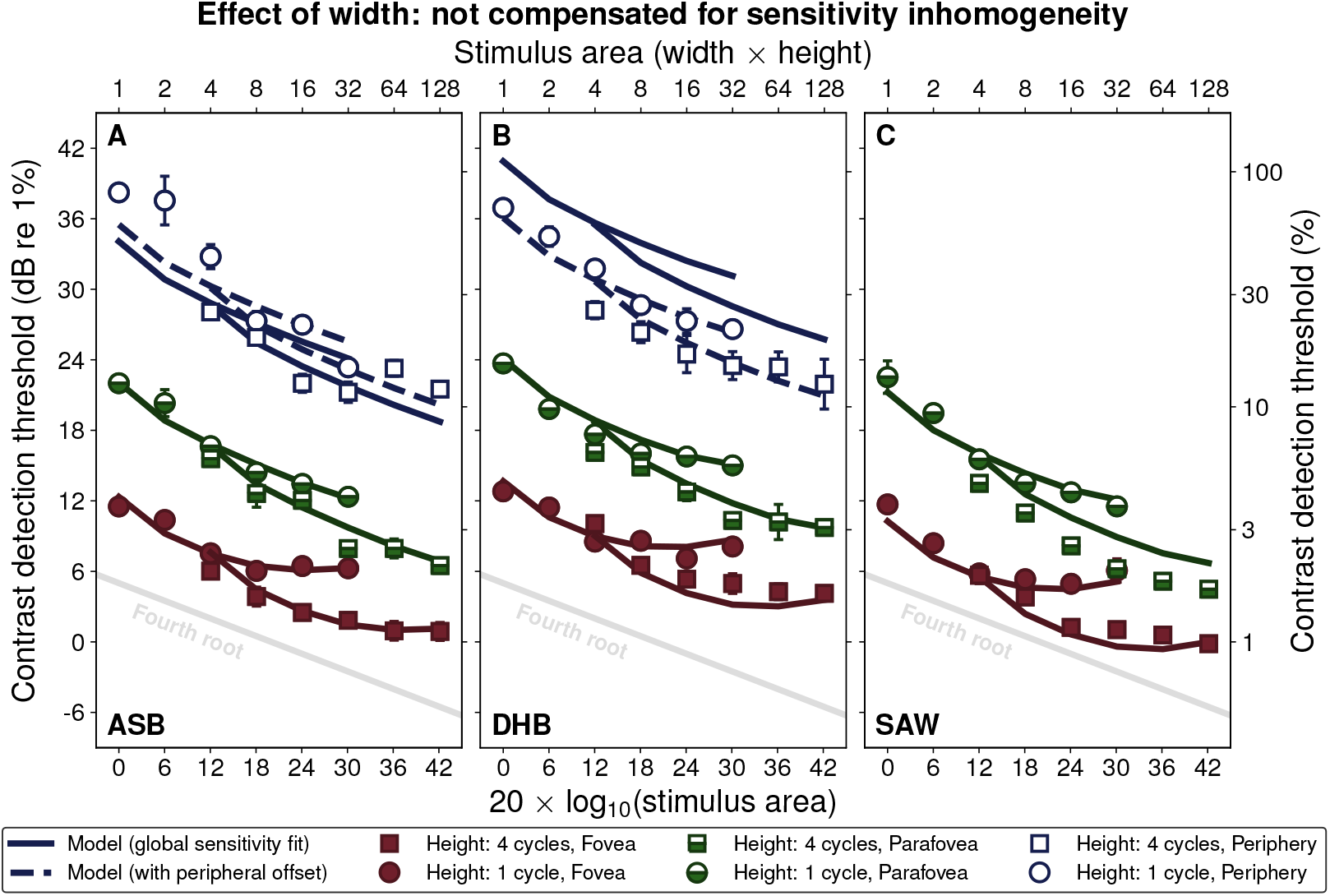
Showing area summation for stimuli with increasing width. The subplots present data from three participants for uncompensated stimuli presented in the periphery (open), parafovea (half-filled), and fovea (filled symbols). Results are shown for stimuli that are 1 cycle (circles) and 4 cycles (squares) in height. The solid and dashed lines are threshold predictions from our model. The grey line shows a “fourth root” summation slope.

In the parafovea and periphery (half-open and open symbols in Figure 3), the benefit from increasing the size of the stimulus extended to larger stimulus areas; the decline in threshold does not flatten out to the same extent as in the foveal condition. This is consistent with previous investigations of summation away from the fovea where stimuli grow along a circular arc centred on fixation (maintaining the same eccentricity) or at a tangent to such an arc (maintaining a *similar* eccentricity). This arrangement largely avoids the confounding effects of the variation in sensitivity across the visual field (e.g. Robson and Graham, 1981; Mayer and Tyler, 1986; Manahilov et al., 2001).

### 3.2 Width summation with Witch Hat compensation

Applying the Witch Hat compensation to our stimuli counteracts the inhomogeneity in local contrast sensitivity, causing summation slopes to “straighten” (Figure 4). This reveals a greater consistency of contrast summation (approximately fourth root) spreading out from the fovea than was apparent with the non-compensated stimuli (those data are replotted with pale symbols in Figure 4 for comparison). We have previously demonstrated a similar result with circular grating stimuli (Baldwin and Meese, 2015). Summation had much the same form across the full 32 cycles of the 4 c/deg stimuli, equivalent to 8 degrees of visual angle.

**Fig 4:**
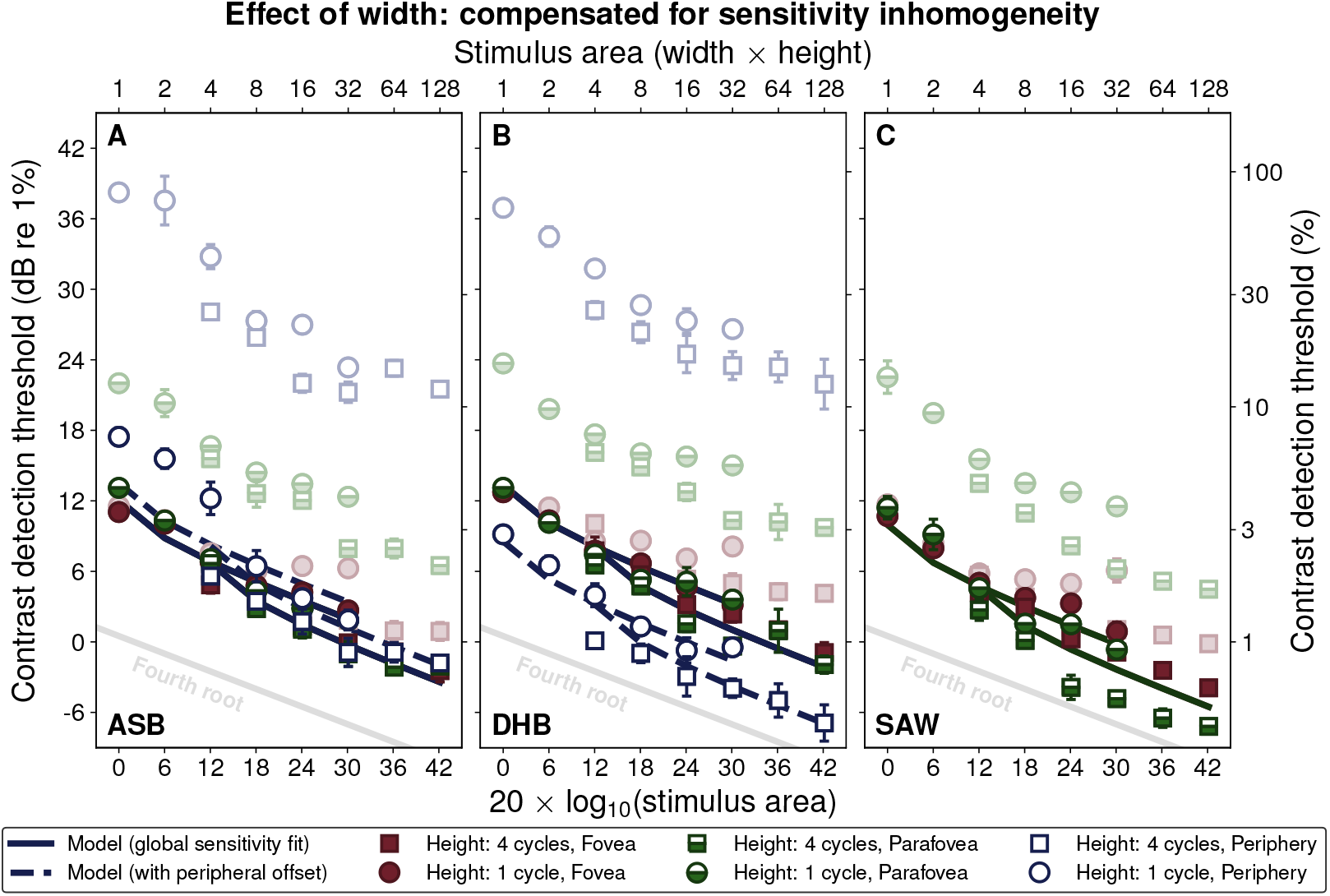
Results for comparable conditions to those in Figure 3, but where local stimulus contrast was adjusted to compensate for visual field inhomogeneity. Results for the uncompensated condition (from Figure 3) are replotted here at reduced contrast. As compensation leads to the overlapping of results from the different visual field locations, we present results separately across field location in Figure 7.

The Witch Hat compensation had less effect on the summation curves for the parafoveal and peripheral stimuli due to the relative homogeneity of contrast sensitivity in those regions. The results for both stimulus heights are consistent with approximately fourth root summation over 32 cycles in the parafovea, a slope shallower than that found by Manahilov et al. (2001) at this eccentricity.

Applying the Witch Hat compensation tended to equate thresholds obtained in the fovea and parafovea for all three participants. In Figure 4, this caused the half-filled (parafovea) symbols to overlap the filled symbols (fovea). This suggests that i) the attenuation surfaces we applied to our stimuli were accurate, and ii) summation in the fovea and parafovea follow the same rule. For participants ASB and DHB, this superposition was striking (Figure 4A-B). For SAW, there was some disparity between the results across the two field positions, the benefit of area for the 4 cycle high stimuli being greater in the parafovea than in the fovea (compare half open squares with filled squares in Figure 4C). We shall return to this point in the **Discussion**.

In the peripheral condition, the stimuli were presented to regions beyond where the attenuation surfaces were mapped in Baldwin et al. (2012). In that study, we measured eccentricities up to 18 cycles, less than half of the distance to the peripheral stimuli in the study here (42 cycles from the fovea at their closest point). We therefore relied on extrapolating the second (shallower) limb of the bilinear decline from our Witch Hat model into the periphery. Nevertheless, the compensation for ASB did a good job of equating sensitivity (the open symbols tend to superimpose with the other symbols in Figure 4A), though there was a tendency for the smaller 1 cycle high stimuli to have higher thresholds than expected. For DHB (Figure 4B) the peripheral compensation overshot, resulting in lower nominal thresholds than for other stimuli of the same size. This implies a shortcoming from our simple extrapolation. In fact, additional measurements at greater eccentricities in Baldwin (2013a) suggest that the second limb of the Witch Hat surface might be slightly shallower than estimated in Baldwin et al. (2012), consistent with the over-compensation for DHB here.

### 3.3 Height summation without compensation for visual field inhomogeneity

Results for the narrowest and widest stimuli from Figure 3 are replotted in Figure 5, with the addition of results for a stimulus height of 2 cycles. These plots show the effect of increasing the height of our narrowest and widest stimuli (triplets of data points). In comparison to Figure 3, where summation that was steeper than “fourth root” was found only for the smallest sizes, we see a greater tendency for summation to outpace the fourth root rule when stimuli grow along the axis parallel to their stripes.

**Fig 5:**
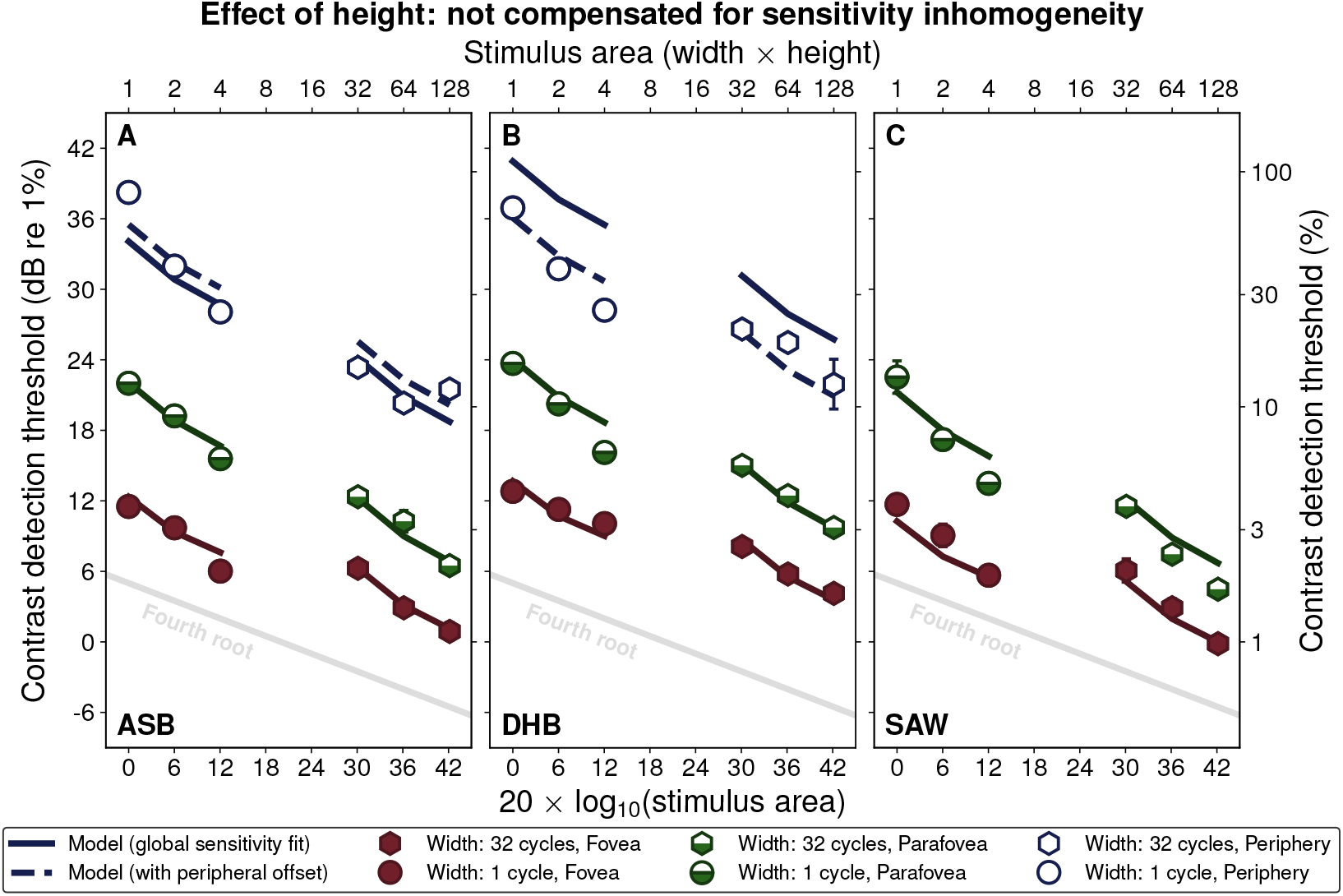
Area summation for stimuli increasing in height (see Figure 3 caption for details on the presentation). Results are shown for stimuli that are 1 cycle (circles) and 32 cycles (hexagons) in width. The first and last datapoint of each triplet (heights of 1 and 4 cycles) are replotted from Figure 3. Intermediate points in each triplet are for a height of 2 cycles.

### 3.4 Height summation with Witch Hat compensation

With Witch Hat compensation applied to our stimuli of different heights (Figure 6), the results from the three visual field locations collapse as before (Figure 4). This also results in similar summation slopes in the fovea, parafovea, and periphery, though the effect of overcompensation from the extrapolated attenuation surfaces is seen again for DHB (Figure 6B; open symbols).

**Fig 6:**
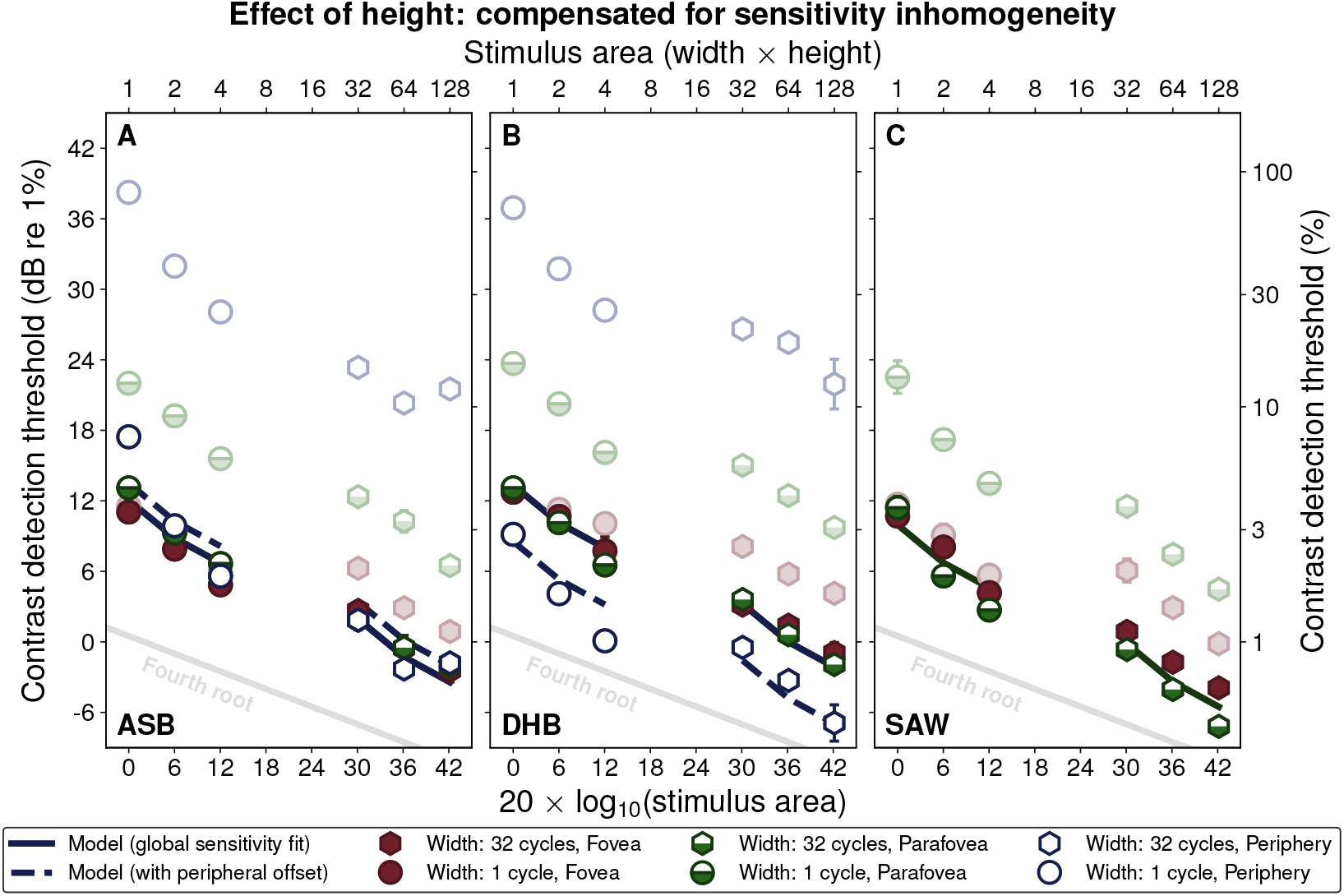
Area summation for stimuli of different heights where local contrast was adjusted to compensate for visual field inhomogeneity (see Figure 5 caption for further details).

### 3.5 Modelling our results

The results in Figures 3-6 are shown with the predictions of the noisy energy model (Meese, 2010; Meese and Summers, 2007, 2012) with the form used in Baldwin and Meese (2015). The model predicts changes in detection thresholds for stimuli of different sizes and locations by finding the contrast that would result in a specific signal-to-noise ratio (*d’* = 1).

To calculate the numerator of the signal-to-noise ratio (the “signal” magnitude for a specific stimulus condition), the stimuli are first attenuated by the Witch Hat surface that was mapped previously for each participant (Baldwin et al., 2012). They are then filtered by Cartesian-separable log-Gabor wavelets (Baker et al., 2022) matched in spatial frequency and orientation to the stimuli. As filters, these wavelets have a spatial frequency bandwidth of 1.6 octaves (full-width at half-maximum) and orientation bandwidth of 25^*°*^. These bandwidths model the responses of neurones with receptive fields analogous to those of simple cells in primary visual cortex (De Valois et al., 1982), with similar values being used in previous models of psychophysical data (Meese, 2010; Schütt and Wichmann, 2017). Within the footprint of these filter elements, responses sum linearly.

The outputs of the local filter-elements are then squared (Meese, 2010) and summed within a template that is matched in size to the outline of the stimuli. The template is not matched to the decline in contrast sensitivity across the visual field. For non-compensated stimuli, the ideal observer would attribute more weight to the regions of the visual field where the sensitivity is highest. However, this detail makes little difference to the model predictions for the stimulus and task design in the current study (Baldwin, 2013b). The square-law transduction of local contrasts in the model has the effect of reducing the summation slope to square root beyond the short-range of linear summation that occurs “within-filter”.

The denominator of the signal-to-noise ratio is the standard deviation of the combined noise affecting the decision. We assume this originates from independent additive Gaussian noise sources affecting the responses of each local filter-element. The larger template sizes include a greater number of these noise sources, with the standard deviation of the combined noise at the decision stage increasing with the square root of the template area. This approach assumes that the observer restricts the template size to the stimulus size within an experimental block, thereby excluding the noise from irrelevant signal locations. By comparison to a fixed summation window (matched to the largest stimulus) this ideal approach (e.g. Tyler and Chen, 2000) reduces the model summation slope from linear to square root. However, with our square-law transduction of signal in place (see above) the combined effects further reduce the model’s summation slope from square root to fourth root.

To fit the model to foveal and parafoveal results, we adjusted a single global sensitivity parameter (an offset) for each participant (i.e., the model has one degree of freedom for each participant). This parameter is equivalent to scaling the common standard deviation of local additive noise sources affecting the response of each filter element.

In our data analysis, we fitted Weibull psychometric functions to find “thresholds” for 82% correct, whereas the model predicted thresholds for a *d*^*′*^ of unity (76% correct in our two-interval forced-choice task). This difference is fully absorbed by our global sensitivity parameter as, for the conditions tested, our noisy energy model produces the same pattern of summation regardless of the criterion *d*^*′*^ used (varying only in the absolute threshold level). Consistent with this, several previous studies have found the slope of the psychometric function to be invariant with stimulus area (including Mayer and Tyler, 1986; Meese and Summers, 2012; Wallis et al., 2013; Baldwin and Meese, 2015).

For each participant, fits were made to all of the foveal and parafoveal thresholds using the *fminsearch* method in Matlab 2016a (Mathworks, Natick, MA). The fitting procedure found the parameter value that produced the lowest RMS error between the model predictions and the human data. The fitted parameter values and RMS errors are shown in Table 1.

**Table 7:** Global sensitivity parameters and RMS errors for the model fits to the results presented in Figures 3-7. A global sensitivity parameter fits the data measured in the fovea and parafovea as described in the text. The offset parameter in the periphery is a second free parameter to adjust sensitivity to the peripheral stimuli alone.

| Participant | Fovea and parafovea |  | Periphery |  |
| --- | --- | --- | --- | --- |
|  | Global sensitivity | RMSe | Offset | RMSe |
| ASB | 11.3 dB | 0.8 dB | 1.4 dB | 2.4 dB |
| DHB | 12.6 dB | 0.9 dB | -4.8 dB | 1.2 dB |
| SAW | 9.2 dB | 1.5 dB |  |  |

Referring back to the model curves for the foveal and parafoveal locations in Figure 3 (solid black lines), the model (with a single fitted parameter) provides a good account of the results for all three participants. It predicts the initial steep (square root) fall in threshold where summation is accelerated by within-filter linear effects, the subsequent fourth root summation, and the differences in sensitivity between the fovea and the parafovea. The model also captures the shallowing of the summation slope caused by the visual field inhomogeneity for the non-compensated stimuli.

In Figure 5, we see how the same model predicts the steeper summation slopes found by growing our stimuli in height (parallel to their stripes) with no further parameters. This may be a simple consequence of linear summation within the simulated “receptive fields” in our model (Meese, 2010), which are elongated at an orientation parallel to the grating stripes. However, it is notable that effects of summation along stimuli with a width of one cycle in the parafovea or periphery outpaces the predictions of our model (which includes this within-filter linear summation) in all three participants. This may point to the influence of specialised mechanisms involved in this type of summation (Chen and Tyler, 1999; Chen et al., 2019, 2023).

For the Witch Hat compensated stimuli in Figure 4, the same model (with the same global sensitivity parameter) predicts that the results for the different field positions should be identical, because the effects of the visual field inhomogeneity have been factored out. The data and model fits are replotted separately for each field position in Figure 7 to facilitate visual comparisons. The empirical results from the fovea and parafovea agree with this prediction with one exception: Participant SAW (Figure 4C) had lower thresholds for the 4 cycle high stimuli in the parafovea than predicted. We consider possible reasons for this in the **Discussion** below.

**Fig 7:**
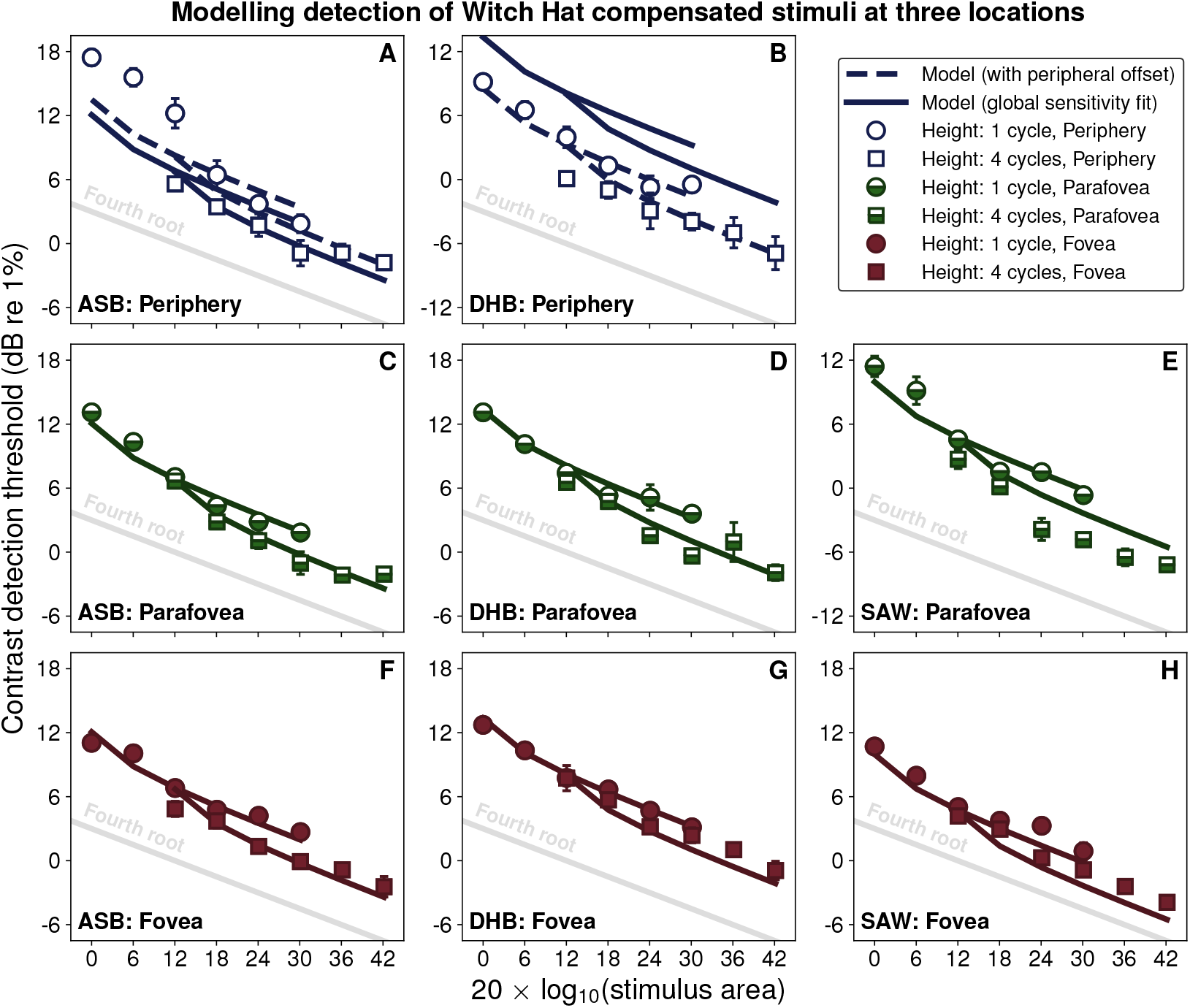
Results for Witch Hat compensated stimuli replotted from Figure 4. Separate panels are for the different field conditions to facilitate comparisons between human data and model predictions at each eccentricity.

Results from two participants (ASB and DHB) for the peripheral condition are also shown in Figures 3-7. Within this region, contrast sensitivity is relatively constant (Robson and Graham, 1981), and so Witch Hat compensation has little effect on the shape of the summation slope. In general, both participants showed fourth root summation as the stimulus size increased up to its maximum width of 32 cycles, consistent with the results reported by Robson and Graham (1981) for this location. However, as mentioned above, the Witch Hat compensation we applied in the periphery is a long extrapolation from the surfaces measured in Baldwin et al. (2012). For ASB, this is quite successful, nonetheless, while for DHB our extrapolated attenuation surface predicted sensitivity to be slightly worse in the periphery than was found. To account for this, we re-fitted these data with an additional “offset” parameter (see Table 1). For ASB the optimal fit required an additional offset of only 1.4 dB since the initial extrapolation was quite close. For DHB rather more correction was required (−4.8 dB).

A final discrepancy between human and model is for ASB’s smallest stimuli. Both with (Figure 7) and without compensation (Figure 3), thresholds were higher than predicted in the periphery for stimuli that were one cycle high and up to four cycles wide. We propose an explanation for this in the **Discussion** below.

## 4 Discussion

### 4.1 Fourth root summation of contrast to threshold across the visual field

We have shown that the spatial summation of contrast to detection threshold can be explained by a common process in the fovea, parafovea, and periphery. Our results are consistent with short-range linear summation within the receptive fields of local detecting mechanisms, followed by fourth root summation across them. We were able to reveal this equivalence by applying our Witch Hat attenuation surface (Baldwin et al., 2012), both as a component in our model and in the rendering of stimuli to compensate for local variations in contrast sensitivity.

In the periphery, our finding that summation for stimuli of more than a few cycles follows a fourth root rule agrees with several previous studies (Robson and Graham, 1981; Mayer and Tyler, 1986, referring to the shallower slope for the larger sizes in the latter study). For smaller stimuli (one to two cycles), we see the effects of linear summation within the model’s filter elements (and presumably the human participant’s early receptive fields). The greater influence of this linear process for the smaller sizes steepens the summation slope beyond fourth root. These steeper slopes are consistent with other studies that tested in the periphery (Manahilov et al., 2001; Meese and Hess, 2007). However, the stimuli in those studies were not as small as those in which we find the steeper slopes in our current study. The noisy energy model can account for this if the design of those studies did not allow participants to restrict their template to the extent of the stimulus. This would defeat one of the two components that combine to give fourth root summation behaviour, resulting in the quadratic summation found in those two studies. For example, in Manahilov et al. (2001) each block was preceded by a suprathreshold example stimulus to reduce uncertainty. The stimuli in that study had a Gaussian spatial envelope, meaning that the “useful” region of the stimulus for detection at contrast threshold would be much smaller than the extent visible in a higher-contrast presentation. This could frustrate the participant’s ability to apply an appropriate template.

In the fovea, although some studies have previously reported fourth root summation slopes (Robson and Graham, 1981; Polat and Norcia, 1998; Polat and Tyler, 1999; Meese and Hess, 2007) in most cases the slopes became shallower as the stimulus size increased. This is expected due to the stimuli growing into regions of the visual field which are less sensitive. Compensating for this inhomogeneity using our Witch Hat model allows us to demonstrate that spatial summation in the fovea, the parafovea, and the periphery follows the same fourth root rule.

One limitation of the current study is that we restricted our testing to the upper visual field, with our stimuli centred on the vertical meridian. This raises the question of whether a common summation rule would also be found in the lower visual field, or for stimuli placed along the horizontal meridian. On the basis that the lower visual field (for example) is likely more similar (in its summation behaviour) to the upper visual field than it is to the fovea, we expect our finding would generalise to other meridians. We previously measured the variation in local contrast sensitivity with the polar angle of the stimulus location (Baldwin et al., 2012), and that effect is incorporated in the Witch Hat attenuation surfaces used in this study. We predict the same fourth root summation rule we found along the upper vertical meridian would persist if we were to instead present compensated stimuli at other locations (within the eccentricity range tested here).

On the other hand, studies that have used different (but related) visual tasks have found effects of polar angle that suggest variations in summation behaviour might occur. For example, research into crowding (a phenomenon in which sets of closely arranged objects, such as letters, presented in peripheral vision seems to be “entangled” in a way that makes it difficult to identify or localise specific members of the set) has found the range over which it occurs to depend on polar angle (Petrov and Meleshkevich, 2011; Greenwood et al., 2017). If crowding arises from the binding of stimulus features involving the same integration processes as in the summation of contrast, then we might also expect differences in summation behaviour with polar angle. Similar effects of polar angle have been reported for a variety of tasks (Himmelberg et al., 2023), including local contrast sensitivity where the variation with polar angle persists even when stimuli are re-scaled by the cortical magnification factor (Jigo et al., 2023). The possibility of a variation in the summation rule with polar angle remains an empirical question that could be tested in future studies.

### 4.2 Modelling the summation of contrast over stimulus area to threshold

The fourth root summation behaviour here is consistent with earlier studies that found evidence to support the noisy energy model (Meese, 2010; Meese and Summers, 2012; Baldwin and Meese, 2015). However, we acknowledge that the results from this study may, on their own, be explained by alternate models that combine spatial filtering with *another* subsequent process that also leads to fourth root behaviour. The main competitor would be the signal detection theory-based probability summation models referenced in the **Introduction** (Tyler and Chen, 2000; Meese and Summers, 2012; Kingdom et al., 2015). However, the results from our other recent work lead us to favour the noisy energy model over a probability summation model (Meese, 2010; Meese and Summers, 2012; Baldwin and Meese, 2015).

Within the domain of additive summation models, the design of our experiments did not allow us to distinguish other possibilities such as a role of contrast gain control (Foley, 1994; Dao et al., 2006; Meese and Summers, 2007) or multiplicative noise at threshold (Lu and Dosher, 2008). The model presented here is relatively simple, having only to predict performance in detecting targets against a blank background. When target stimuli are detected or discriminated at contrasts *above* the detection threshold, we expect inhibitory or “gain control” mechanisms to play a substantial role (e.g. Legge and Foley, 1980; Meese and Baker, 2011; Baldwin et al., 2016). Recent work on summation above threshold contrast in collinear (“skunk tail”) patterns suggests that these inhibitory signals may be pooled similarly (same summation rule and spatial extent) to the excitatory signals that dominate performance at threshold (Chen et al., 2019, 2023).

In general, the noisy energy model provides a very good account of the results across the different stimulus sizes and visual field locations. It did this with only one free parameter (controlling the global sensitivity) per participant for the foveal and parafoveal results, and with one extra parameter (an offset to account for inaccuracies in the extrapolated Witch Hat model) for the peripheral results. There are two cases, however, where individual participants showed consistent idiosyncratic deviations from the model prediction. The first is for participant SAW (Figures 3C and 4C), where thresholds were systematically lower than predictions for stimuli that were four cycles high and presented in the parafovea. One possible explanation is an individual difference for this participant involving elongated receptive fields in the parafovea (Wilson and Sherman, 1976) with more extensive short-range linear summation within filter elements.

The second deviation from our singular model is found in participant ASB, where thresholds for the smaller stimuli (one cycle high and up to four cycles wide) were approximately 3 dB higher in the periphery than predicted (Figures 3A and 4A). If this result had been seen for all stimuli with a height of one cycle, then a similar “elongated receptive field” explanation could be offered as for SAW. Although ASB’s data fell *above* the prediction, whereas SAW’s fell *below* it, this could be due to inaccuracy of the extrapolated Witch Hat (that would then be accounted for by our peripheral offset parameter). Unfortunately, such an explanation does not suffice, as the increased thresholds seen for the one cycle high stimuli presented to ASB in the periphery are not found for *widths* greater than four cycles. A possible explanation can be found in models of summation along “skunk tail” stimuli, which suggest there may be additional, more extended, receptive fields that pool along the stripes of a grating stimulus. Relevant to our result is that this second-order filter stage is proposed to differ in its behaviour in the periphery (e.g. becoming phase-insensitive, where it is phase-sensitive in the fovea, see Chen and Tyler, 1999).

An alternate approach to account for ASB’s anomalous results would be a constraint on the template stage in our model. For example, if the participant was unable to match a template to stimulus area for small peripheral stimuli, then the integration of additional noise in a mismatched template would reduce the signal-to-noise ratio for those stimuli, causing thresholds to lie above the model predictions (as we found). This would not have been compensated by the Witch Hat, because this region was not mapped by Baldwin et al. (2012). Such a limitation on the template stage might result from uncertainty about the location of the stimulus in the periphery, or mandatory signal combination over a minimum summation region (as in crowding, e.g. Parkes et al., 2001). Further work would be required to determine whether this minimum summation region simply corresponds to the second-order filter stage proposed by Chen and Tyler (1999).

Between the measurement of the Witch Hat surface (in Baldwin et al., 2012) and the summation experiments themselves, the duration of testing required for each participant in the current study limited our ability to increase our sample size and go on to investigate individual differences further. However, the current study does motivate the design of future work to investigate systematic variations in summation between individuals. These differences may have profound implications for the mechanistic basis of summation (Mollon et al., 2017).

## 5 Additional information

## 5.1 Acknowledgements

We would like to thank Rob Summers for the Liberator software used to perform this study. This research was supported by an Engineering and Physical Sciences Research Council (EPSRC, UK) grant to TSM and Mark Georgeson (EP/H000038/1), and by a Natural Sciences and Engineering Research Council of Canada (NSERC) Discovery Grant awarded to ASB (RGPIN-2022-04216).

## 5.2 Author contributions in CREDIT format

### ASB

Conceptualization, Data Curation, Formal Analysis, Investigation, Methodology, Software, Visualization, Writing - Original Draft.

### TSM

Conceptualization, Funding Acquisition, Investigation, Methodology, Project Administration, Supervision, Writing - Review & Editing.

## Footnotes

2 A separate line of research concerns summation of broadband luminance spots, discs, or Gaussian blobs, (e.g. Bijl and Koenderink, 1993). Previous work on this type of summation in the periphery is reviewed by Strasburger et al. (2011).

